# Dynamic valuation bias explains social influence on cheating behavior

**DOI:** 10.1101/2024.05.21.594859

**Authors:** Julien Benistant, Valentin Guigon, Alain Nicolas, Edmund Derrington, Jean-Claude Dreher

**Affiliations:** CNRS, Institut des Sciences Cognitives Marc Jeannerod, Bron, France; Univ. Lille, CNRS, IESEG School of Management, UMR 9221 - LEM - Lille Économie Management, F-59000 Lille, France; Université Lyon 1 Claude Bernard, France; Genopsy, CH Vinatier, Bron, France

**Keywords:** social influence, computational model, lPFC, neuroeconomics

## Abstract

Observing immoral behavior increases one’s dishonesty by social influence and learning processes. The neurocomputational mechanisms underlying such moral contagion remain unclear. We tested different mechanistic hypotheses to account for moral contagion. We used model-based fMRI and a new cheating game in which participants were sequentially placed in honest and dishonest social norm contexts. Participants’ cheating behavior increased in the dishonest norm context but was unchanged in the honest. The best model to account for behavior indicated that participants’ valuation was dynamically biased by learning that others had cheated. At the time of choice, the internalization of social norms was implemented in the lateral prefrontal cortex and biased valuations of cheating. During learning, simulation of others’ cheating behavior was encoded in the posterior superior temporal sulcus. Together, these findings provide a mechanistic understanding of how learning about others’ dishonesty biases individuals’ valuation of cheating but does not alter one’s established preferences.

**Significance statement:** Social influence is at the root of human behavior. For example, we tend to follow others’ bad moral behavior such as cheating. Here, we explore the neuro-computational mechanisms of social influence on cheating behavior. We validated a new model capturing both how we learn about others’ (dis)honesty and how this bias our choice. We show that if we observe dishonest others we tend to be more dishonest ourselves. This behavioral change is driven by a bias dynamically changing with our knowledge about the others’ cheating behavior. Neurally, we found that the lateral prefrontal cortex implements this bias into the decision process while the posterior superior temporal sulcus and the temporo-parietal junction encode our learned representation of others’ cheating.

## Introduction

Dishonest behavior, such as cheating, tax evasion and corruption is pervasive in modern societies. Dishonest behavior can be modified through the exposure to others’ immoral behavior (1, 2). For example, observing others’ dishonest reports increase one’s likelihood to cheat (2). Such influence of others’ behavior on our own choices is not limited to dishonesty, but can be observed in domains such as risk-related decision making (3–8), and both pro-social and anti-social behavior (9–12). Why and how the observation of others’ dishonest behavior may modify our own moral behavior in non-strategic settings remains an enduring puzzle. Recent work has implicated specific brain regions in moral decision-making and described the underlying neural computations (13–17). Yet, little is known about the neurocomputational mechanisms underlying how social influence modifies moral decisions.

It remains unclear whether social influence shapes our individual beliefs, as behavior reflecting social norms compliance is not a perfect reflection of our private beliefs. To date, two mechanisms have been proposed to explain how social influence affects one’s choices. The first, called valuation bias, proposes that individuals assign value to social information, which then influences one’s personal valuation process. This means that simply being exposed to others’ choices alters how individuals perceive the value of the options at stake and potentially influence one’s own choice (3, 4, 18). The second mechanism, referred to as preference shifting, proposes that exposure to others’ choices can more fundamentally modify an individual’s own preferences to align with the choices of others. This mechanism implies a more profound change of private beliefs or personal preferences (3–8). One limitation of these two previous accounts of social influence is that they did not consider the dynamic aspect of learning. For example, in a new social environment with specific norms (e.g., when one arrives in a new company), one usually lacks knowledge about others’ behavioral tendencies, including whether they lean toward honesty or dishonesty. Consequently, in such new environments, learning through repeated observation of others’ behavior becomes necessary to mitigate uncertainty about others’ true levels of (dis)honesty. Learning social norms in new contexts is particularly important regarding dishonesty because of its concealed nature (because this makes accurate observation challenging). Therefore, it is likely that a social learning component might modulate either the valuation bias or the preference shifting mechanism, to adjust how such mechanisms are weighted as the individual learns about others’ honesty levels.

Here, we tested different theoretical accounts of the mechanisms underlying how others’ behavior influences one’s own cheating behavior. We considered four social influence mechanisms, as either a fixed or dynamic valuation bias or a fixed or dynamic change in individual preferences (Fig. 1). First, we assessed which computational model among these best explained social influence in cheating behavior. Second, we uncovered where in the brain different computational signals of social influence are encoded when presented with the opportunity to cheat. Importantly, the different accounts noted above predict that different neural mechanisms underly social influence on cheating behavior. According to the first two hypotheses (fixed valuation bias and fixed preference shifting hypotheses), no behavioral or brain activity changes should occur over time, because these hypotheses do not consider that learning about the behavior of others would dynamically affect the influence process. In contrast, the dynamic valuation bias and dynamic preference shifting hypotheses do predict such changes. One brain region that could support such dynamic changes in social influence is the dorsolateral prefrontal cortex (dlPFC) because it represents others’ risk-preferences, and because its functional connectivity with the ventral striatum increases with the extent of social influence (8). Moreover, activity in the dlPFC is associated with integration of social norms and moral preferences in decision processes (16, 19–21) and neuroimaging studies consistently implicate the dlPFC in dishonest choices (22–24). Regions such as the right temporo-parietal junction (rTPJ) and the posterior superior temporal sulcus (pSTS) are also good candidates to support the dynamic integration of others’ behavior. Indeed, both the rTPJ and pSTS are central in tracking and integrating others’ preferences (25), and there is evidence that the rTPJ encodes others’ intentions necessary for social choices (26–29).

**Figure 1:**
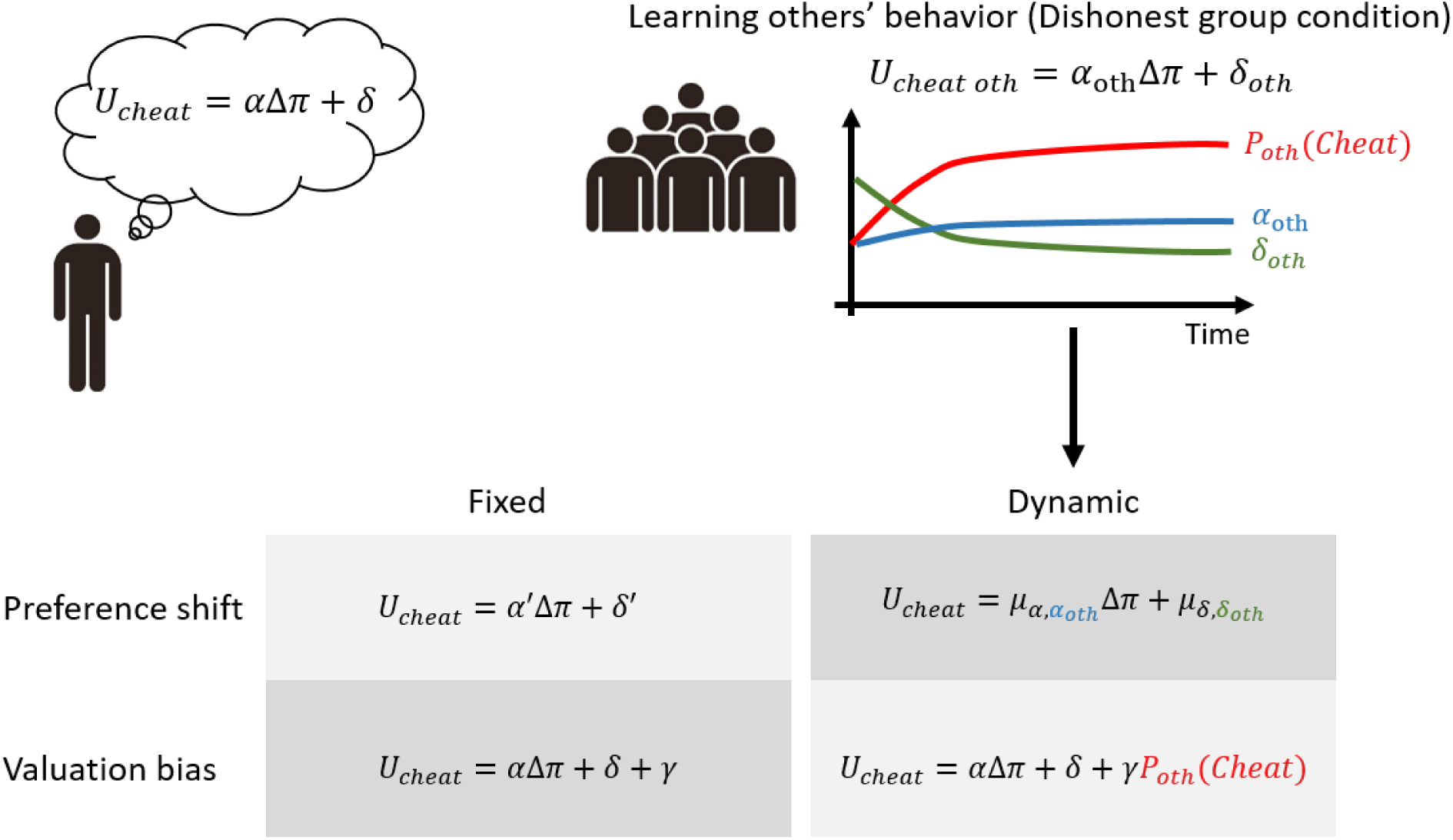
Graphical representation of the different computational models accounting for social influence. This figure represents the 4 different mechanisms of social influence that we considered. We varied whether social influence is due to a shift in one’s preferences (top line of the table) or is due to a bias in the valuation process (bottom line). We then vary whether the process is fixed (left column) or dynamic (right column) because it depends on one’s learning about the behavior of others over time (schematically represented by the top figure, right column). For each model a formal description is provided based on a utility function (U_cheat_) representing the relative value of cheating for the individual, with two free parameters α and δ. α represents the individuals sensitivity to monetary reward whereas δ represents her moral cost faced when cheating.

We used functional magnetic resonance imaging (fMRI) and a new paradigm based on a cheating game. This game comprised two types of trials: (i) Solo trials, in which participants could lie about the outcome of a die colzaroll to maximize their earnings, (ii) Predict trials, in which they predicted what another individual, randomly selected from a group of 10, reported in a previous experimental session. The purpose of the Solo trials was to assess the extent to which participants were prone to cheating. For the Predict trials, the goal was to expose participants to the behavior of others and allow them to learn the preferences of a group. Unbeknownst to the participants, the behavior of the group was simulated and controlled so that the participant faced either a dishonest or an honest group of players. With this manipulation, we assessed the effect of social influence in two different contexts. These two contexts have been shown to lead to different levels of conformity because anti-social and dishonest behavior appears to be more contagious than prosocial behavior (11, 30, 31). The experiment was divided into three blocks. The first was composed of trials, to allow us to estimate participants’ preferences in the absence of social influence. The second and third blocks consisted of interleaved Predict and Solo trials. In each of these two blocks, participants’ predictions concerned either an honest or a dishonest group of participants. This novel design gradually exposed participants to others’ behavior, allowing us to test the fixed *versus* dynamic accounts of valuation bias and preference shift mechanisms.

## Results

### Experimental design

We scanned 32 participants using fMRI while they played the cheating game, which included two types of trials: Solo and Predict (Fig. 2.A). Solo trials required participants to observe a die roll result (‘stimulus’). Then, participants were presented with two dice (‘choices’), one presenting the original die roll result and the other showing an inaccurate die roll. Each die was associated with a different payoff. Notably, the accurate die always offered a lower payoff than the inaccurate one, which presented participants with a moral dilemma between honesty and maximization of their earnings. To modulate the difficulty of these dilemmas, the payoffs varied across trials. Participants had as much time as they wanted to choose which die roll to report. Once their choice was made, the screen froze for 500 ms (‘confirmation’). No visual feedback indicating their choice was provided to reinforce the feeling that the decision was made privately (see SI Methods for details).

**Figure 2:**
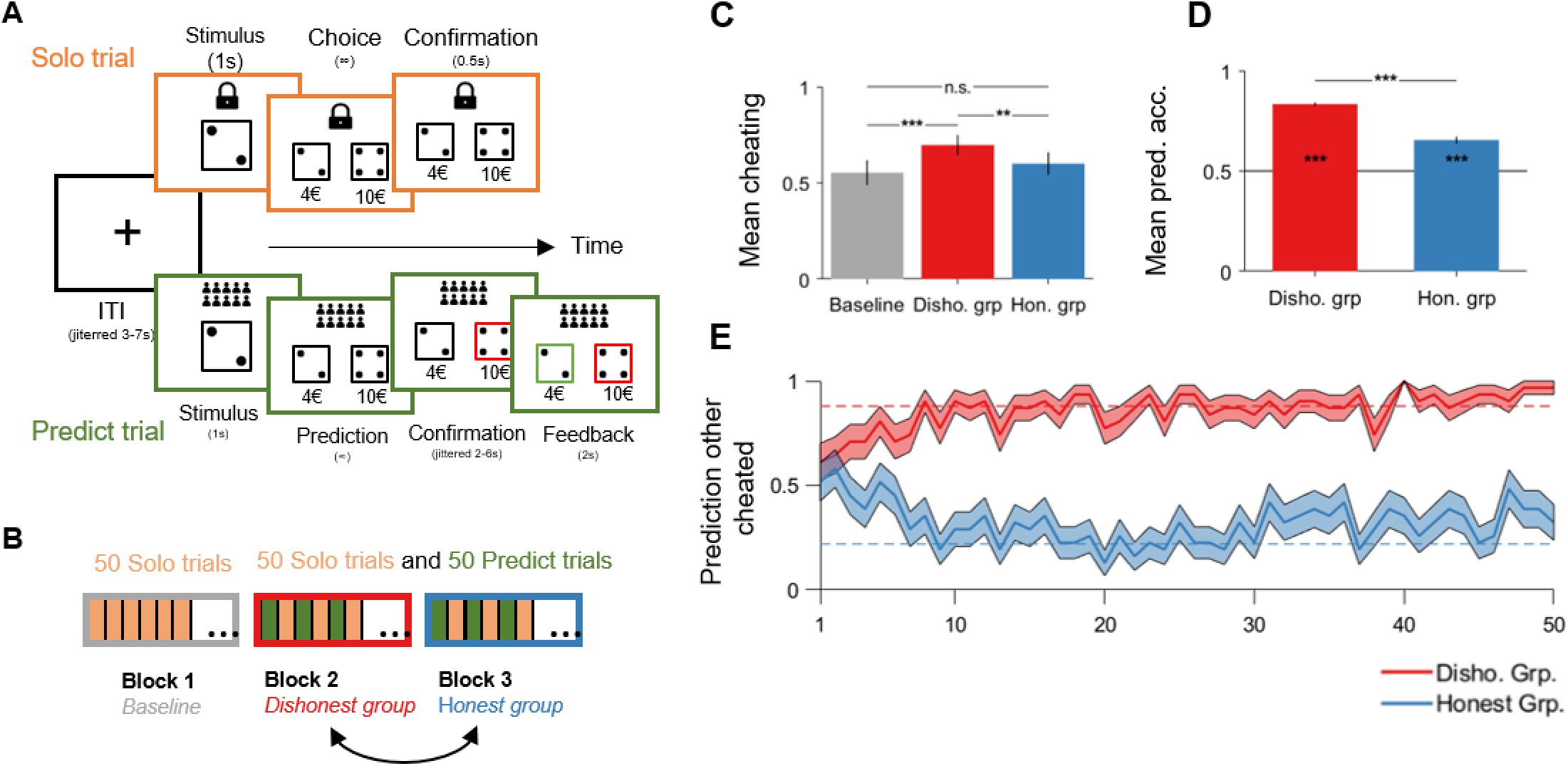
Experimental design and behavioral results. **A.** The experiment consisted of 2 types of trials. In the Solo trials participants had to report the outcome of a die draw among two choices. An honest report was always less rewarded than a dishonest one making cheating the more profitable choice. The earnings associated with each choice varied between trials and ranged from €0 to €8 (€2 to €10) by steps of €2 for the honest (dishonest) choice. The Solo trials ended with a confirmation step in which the participants’ screen froze without confirmation of her choice to reinforce the feeling that decision was anonymous. In the Predict trials, participants had to predict whether another individual, randomly selected from group of 10, cheated or reported honestly. Then, the participants’ response was highlighted in red during a jittered confirmation step. Finally, participants received feedback about their prediction. The correct response was highlighted in green. **B.** Our experiment was divided in 3 blocks. The first one, the Baseline, consisted of 50 Solo trials and allowed us to assess the participants’ preferences. The two following blocks were a mixture of 50 Predict and 50 Solo trials that were interleaved. This structure allowed us to expose participants to the others’ behavior and to assess the effect on participants’ decisions. In each of these two blocks, participants’ predictions concerned either an honest or a dishonest group of participants. The order of presentation of the Honest and Dishonest Groups was randomized between participants as either the second or third blocks. **C**. The mean proportion of cheating in the Dishonest Group condition was significantly higher than in the two other conditions. The statistical analysis comes from a pairwise comparison with Bonferroni correction extracted from a mixed-effect logistic regression (see model 1 in Table S2 for details). **D**. Mean prediction accuracy in both the Dishonest Group and the Honest Group conditions. The black line indicates chance level. Stars within bars indicate the results of a one-sided signed rank test comparing the prediction accuracy with the chance level. The other statistical analysis comes from a mixed-effect logistic regression (see model 2 in Table S2 for details). **E**. The mean of predictions that the other cheated over Predict trials for the both the Dishonest and Honest Group conditions. The red (blue) dashed line corresponds to the mean proportion of cheating of the Dishonest (Honest) group members. In all figures, bars or shaded areas corresponds to standard errors. *** p < 0.001, ** p < 0.01, * p < 0.05.

The “Predict” trials mirrored the Solo trials but required participants to predict the choice of whether another individual would report their own die roll accurately or not in similar circumstances. Participants were informed that this individual was randomly chosen from a pool of 10 anonymous individuals that had completed the task previously. In reality, this group was computer simulated, which allowed controlled manipulation of their cheating behavior. The prediction of the participants was highlighted in red during a jittered ‘confirmation’ stage. Finally, at the end of each Predict trial feedback was provided to the participant by highlighting the correct response in green. Accurate predictions were rewarded, whereas inaccurate predictions were not (see SI Methods for details).

The experiment comprised three blocks: a Baseline block with 50 Solo trials to discern individual preferences in the absence of social influence, and two subsequent blocks each consisting of 50 Solo and 50 Predict trials interleaved (Fig. 2.B). This sequential arrangement aimed to gradually expose participants to others’ behavior, to test how learning about the others’ (dis)honesty influenced their own decisions in the Solo trials. In these two blocks, one block featured dishonest others (Dishonest Group condition), while the other had honest counterparts (Honest Group condition, see SI Methods and Table S.1). The presentation order of the latter two blocks was randomized across participants.

From the initial pool of 32 participants, one was excluded from all analyses due to consistent cheating and disbelief regarding whether they were predicting real peoples’ decisions. Additionally, three more participants were excluded from the model-based analyses because they always cheated or always were honest in the Baseline Solo trials (see SI Methods).

### Behavioral effect of social influence

In the Baseline participants cheated in 55.3 % of the Solo trials. They cheated in 69.6 % of Solo trials in the Dishonest Group condition and in 60 % of the Solo trials in the Honest Group condition (Fig. 2.C). To test the differences between each condition, we used a mixed-effect logistic regression on whether the participants cheated or not in a given trial with the following independent variables: a categorical variable coding for the condition (1: Baseline, 2: Dishonest Group and 3: Honest Group); a binomial variable coding for the order of presentation of the Dishonest Group condition (1: first and 0: second); a variable coding for the trial number within each block; a variable coding for the difference between the payoff of cheating and honesty; and, a variable coding for the absolute difference between the die value of the cheating option and the honest one. We also added demographic variables such as the participants’ sex, age and occupation (*i.e.,* student or not). All the regressions reported in this paper used the same independent variables except when indicated otherwise.

Participants were significantly more likely to cheat in the Dishonest Group condition than in any other condition (Fig. 2.C, margins contrast Dishonest *>* Baseline: 0.148 ± 0.037, *p <* 0.001; margins contrast Dishonest *>* Honest: 0.101 ± 0.027, *p* = 0.001; see model (1) in Table S2). No significant difference in cheating behavior was observed between the Honest Group condition and the Baseline (Margins Honest *>* Baseline: 0.047 ± 0.025, *p* = 0.170). To ensure that these differences were driven by the groups’ behavior, we tested whether participants learned equally well in both conditions. This was indeed the case because participants’ prediction accuracy was significantly different from chance level (i.e., 50%) for both groups (Signed rank tests, *p <* 0.001 for both groups; Fig. 2.D). Fig. 2.E shows that, on average, participants’ predictions about others’ cheating frequency converged towards the mean cheating behavior in both conditions. Moreover, predictions that others would cheat were significantly more frequent for the Dishonest Group condition than the Honest Group condition (Margins Dishonest *>* Honest: 0.540 ± 0.033, *p <* 0.001; see model (3) in Table S2). However, participants were also significantly more accurate when predicting the Dishonest group’s behavior than that of the Honest group (Margins Dishonest *>* Honest: 0.179 ± 0.022, *p <* 0.001; see model (2) in Table S2).

### Computational models of social influence

Our computational approach to analyze social influence encompassed two components: a social learning component and a social influence component. The social learning component accounts for how participants predicted others’ cheating behavior and updated their own beliefs about others’ preferences. The social influence component reflects how participants cheated (or not), based on their own preferences and based upon how they were influenced by what they learned about others’ cheating behavior. To simplify the selection process between our candidate models, we used a three-step selection procedure based on a group-level random-effect Bayesian model selection (32). We started by defining which utility function, among the fixed moral cost and the variable moral cost, best described how participants chose between the honest and the dishonest option. Subsequently, we determined the best fitting learning model that explained participants’ predictions in blocks 2 and 3. Finally, we conducted model selection among the four candidate models that represent the different social influence mechanisms. Below, we describe each of these steps in more detail.

First, we used the participants’ decision to cheat (choose the Dishonest option) in all three blocks. Previous work shows that honesty-based choices can be described by two utility functions. A first utility function (Fixed cost) assumes that individuals endure a fixed moral cost when they are cheating (33, 34). A second utility function (Variable cost) assumes that the cost of being dishonest is dependent on the absolute gains associated with it (24) (see Methods SI). We considered extreme cases in which participants always used just one of these two utility functions (Fixed cost only or Variable cost only) and others cases in which they used one in some blocks and the other in the other blocks (Mixed variants). Results from the model selection showed that the utility function with a fixed moral cost of cheating best explained participants’ cheating behavior across all blocks (Protected exceedance probability (pEP) = 0.999, Fig. S1.A).

Next, we evaluated which model best explained participants’ social learning behavior during the Predict trials in the Dishonest and Honest Group conditions. We tested various candidate models derived from the Bayesian Preference Learning (BPL) model (35). These models initiate with participants holding normally distributed priors regarding others’ preferences, which allow participants to infer the likelihood of cheating when they predict others’ choices during the Predict trials. Post-feedback, a prediction error, that compares their prediction with the actual behavior, refines the posterior distribution of others’ preferences. Within the BPL model, we considered three types of priors, each representing distinct learning biases. The first involved participants using their own preferences as priors for both conditions, a process termed self-projection (BPL-Self). The second utilized participants’ priors based on their own preferences for the initial group condition and the learned preferences from the preceding group for the subsequent group condition (BPL-Others). This showcases the persistence of prior learning. Finally, the third model combined aspects of the previous two types of priors. When learning about the second group’s behavior, participants combined their own preferences with the learned preferences regarding the previous group (BPL-SelfOthers). Furthermore, we investigated whether participants inferred others’ preferences according to the utility functions previously mentioned. Specifically, participants could believe that others’ decisions aligned with the utility function used to simulate others’ behavior (Variable cost of cheating, (24)), or they might consider the utility function they used for their own decisions (Fixed cost of cheating) ((33, 34); see Methods SI). We found that the BPL model with self-projection (BPL-Self) and the utility function with a fixed moral cost of cheating was the best to explain our participants’ predictions (pEP: 0.907, Fig. S1.B).

Finally, we proceeded to the selection of the model that best explains social influence in our experiment. We explored two hypotheses, each delineating social influence in distinct ways: one as a fixed phenomenon, independent of participants’ learning about others’ cheating behavior; and the other as a dynamic and learned phenomenon, interconnected with participants’ understanding of others’ conduct.

For the fixed hypothesis, we evaluated two models. The first posits that social influence arises from a shift in individuals’ preferences (Preferences Shift, PS-Fixed) (5, 7, 8, 12). The second suggests that social influence results from a bias in the valuation process (Valuation Bias, VB-Fixed) (4, 18). For the dynamic hypothesis, we examined versions of these models in which influence correlated with what participants learned of others’ cheating tendencies across trials. Thus, in the PS-Dynamic model, preference alteration was hypothesized to result from a weighted average between participants’ own preferences and the inferred preferences of others at a given time (PS-Dynamic model). This weighted average is governed by two distinct free parameters, 𝛾𝐴 and 𝛾𝐷, for each parameter of the others’ utility function (𝛾𝐴 for the parameter α and 𝛾𝐷 for the parameter δ). They are specific to each group condition, indicating the participants’ levels of conformity. For the VB-Dynamic model, we considered that the participants’ decision model’s value is modulated by the probability that others would have cheated (or been honest) in the Dishonest Group (Honest Group) condition during a particular Solo trial, as calculated by the BPL Self model (VB-Dynamic model). A parameter, 𝛾, adjusts the probability and represents the extent of participants’ conformity to others’ behavior. This parameter was individually estimated for each participant in each group condition.

In total, we tested 4 models (see Fig. 1 for a graphical summary). The Bayesian model selection showed that the VB-Dynamic model was the most frequent best fit across our sample (pEP = 0.672; Fig. 3.A and Figure S2 for a graphical representation of the model).

**Figure 3:**
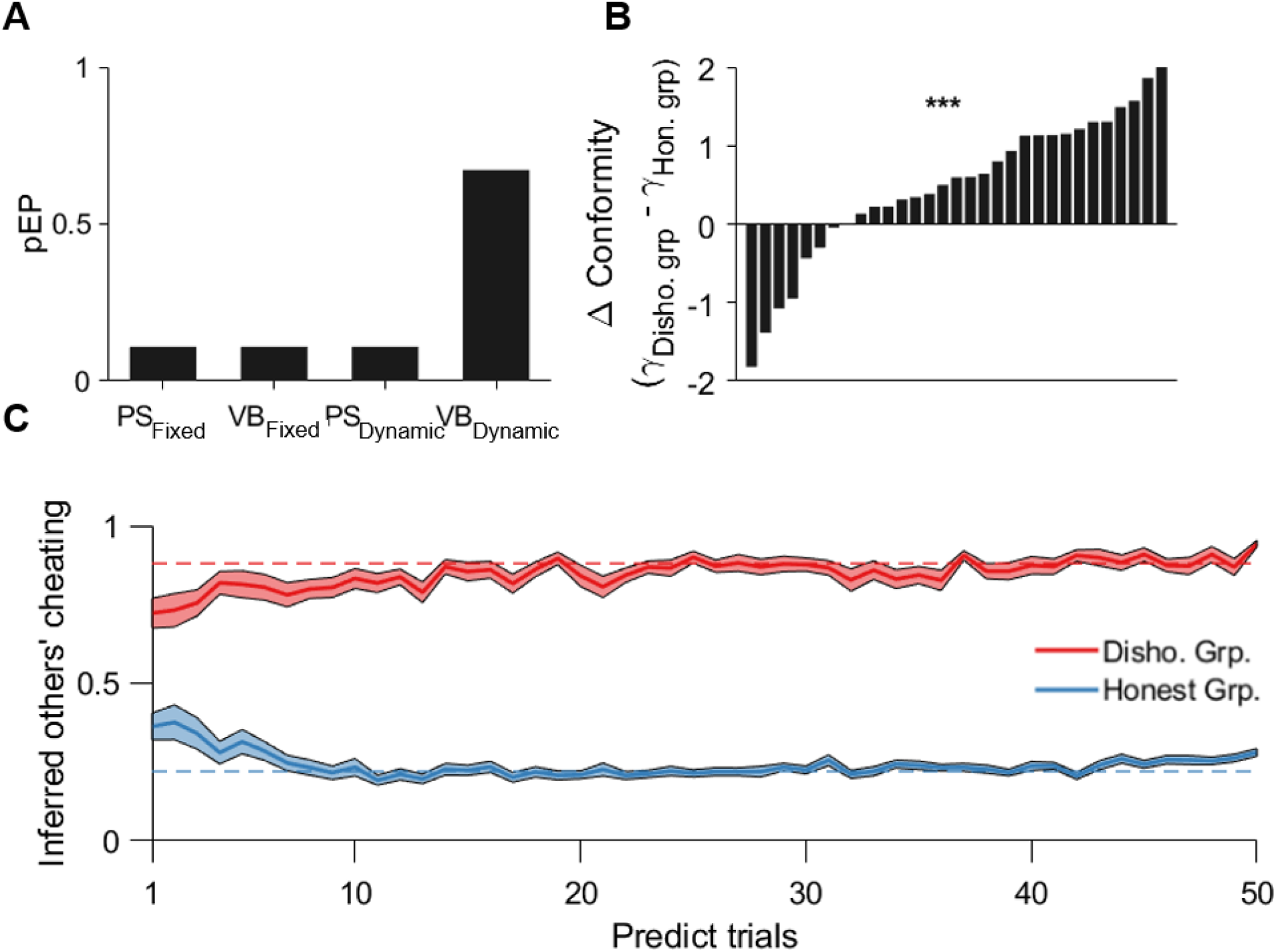
Model selection and estimation. **A.** Protected Exceedance Probability (pEP) of the different candidate models. The higher the pEP, the more likely a given model explained the group’s behavior more than the others. PS: Preferences shift; VB: Valuation Bias. **B.** Difference between the value of the conformity parameter 𝛾 in the Dishonest Group condition and the Honest Group condition for each participant. The stars represent the result of a two-sided rank sum test between the two conditions. **C.** Mean inferred probability that others cheated over Predict trials for the Dishonest Group condition (red) and the Honest Group condition (blue). Shaded areas are standard errors. The red (blue) dashed line corresponds to the mean proportion of cheating of the Dishonest (Honest) group members. *** *p <* 0.001.

We then explored the estimated parameters of the winning VB-Dynamic model. First, we observed that the conformity parameter 𝛾 is significantly higher in the Dishonest Group condition than in the Honest Group condition (Two-sided rank sum test, p<0.001; Fig. 3.B). The decision model parameter *α* is positive and significantly different from 0 (mean: 0.192 ± 0.022, *p <* 0.001, Fig. S2.A) whereas *δ* is negative and significantly different from 0 (mean: −0.684 ± 0.178, *p* = 0.001, Fig. S2.A). This is in line with the fact that participants’ likelihood to cheat increased with the relative earnings of cheating and that this effect was limited by a fixed moral cost associated with cheating. Concerning the learning of others’ preferences, we found that the predicted probability to cheat of others, estimated by participants, converged towards the average simulated probability to cheat of each group (Fig. 3.C).

Formally, the learned *α* parameters are significantly higher for the Dishonest Group than the Honest Group. Conversely, the learned moral cost *δ* parameters are significantly lower for the Honest Group than the Dishonest Group (mean *α*, Dishonest group: 0.428, Honest Group: 0.102; mean *δ*, Dishonest group: −0.089, Honest Group: −1.893; Two-sided rank sum tests, *p <* 0.001 for each parameter; Fig. S2. B-C). This implies that participants recognized that members of the Dishonest Group were more likely to cheat than members of the Honest Group for a given set of payoffs. The difference observed in prediction accuracy between the two groups is not explained by a difference in inverse temperature *β*. A lower value of this parameter for the Honest Group would indicate more randomness in the participants’ predictions (Two-sided rank sum test, *p* = 0.573; Fig. S2.B). However, the variance parameter of the distribution for the learned parameter *α* is significantly higher for the Honest than the Dishonest Group. Conversely, the variance parameter of the distribution for the learned parameter *δ* is significantly higher for the Dishonest Group than the Honest one (mean var. *α*, Dishonest group: 0.378, Honest Group: 0.774; mean var. *δ*, Dishonest Group: 0.529, Honest Group: 0.359; Two-sided rank sum tests, *p <* 0.001 and *p* = 0.003, respectively). Furthermore, the difference in variance for the parameter *δ* is significantly lower than the difference for the parameter *α* (Two-sided rank sum tests, *p <* 0.001). This finding for the model parameters parallels the behavioral results which indicates that participants’ prediction accuracy is lower in the Honest Group condition than the Dishonest Group condition.

Finally, one could argue that participants’ conformity may be explained by their own preferences or by the accuracy of their predictions. These alternative hypotheses were in fact ruled out by additional analyses showing that 𝛾 is neither correlated with the participants’ own moral cost δ, nor with participants’ mean prediction accuracy (Fig. S2. D-E).

### Model-based fMRI analysis

We constructed two GLMs to identify how the brain encodes the 4 computational signals of the VB-Dynamic model. The first 3 signals concerned the decision and prediction processes (at the time of choice in the Solo trials and the time of prediction in the Predict trials): (1) The dynamic valuation bias in the Dishonest and Honest Group conditions (choices in the Solo trials); (2) the prediction that the others cheated, when observing Dishonest and Honest group members (Predict trials), (3) the relative value of chosen and unchosen options in Solo trials, regardless of conditions (i.e., all blocks averaged together). The fourth signal concerned the prediction error signal at the feedback stage of the Predict trials. Four parametric regressors of no interest were also added: participants’ response times, the side of their choice (left or right), their decision to cheat or not at the choice stage of the Solo trials and their predictions about others’ behavior during the prediction stage of the Predict trials. All the parametric regressors were non-orthogonalized, and could thus compete for variance. Finally, we added the participants’ degree of conformity *γ* for both the Dishonest and Honest Group conditions as a second level covariate.

In the first GLM (GLM 1), we included the first signal (dynamic valuation bias) as well as the participants’ uninfluenced relative decision value to cheat as parametric regressors during the choice phase of the Solo trials in both the Dishonest and Honest Group conditions (see Methods SI). This GLM revealed a negative relationship between left Lateral Prefrontal Cortex (lPFC) activity that corresponded to the dynamic valuation bias signal with participants’ conformity parameter 𝛾, for Solo trials in the Dishonest Group condition only (lPFC, x,y,z = −39,27,9; Fig. 4.A-B; p*<*0.05 whole-brain cluster corrected family-wise error (FWE) and Table S3). That is, for negative values of 𝛾, that corresponded to anti-conformity, participants’ dynamic valuation bias signal increased more in the lPFC at the time of choice.

**Figure 4:**
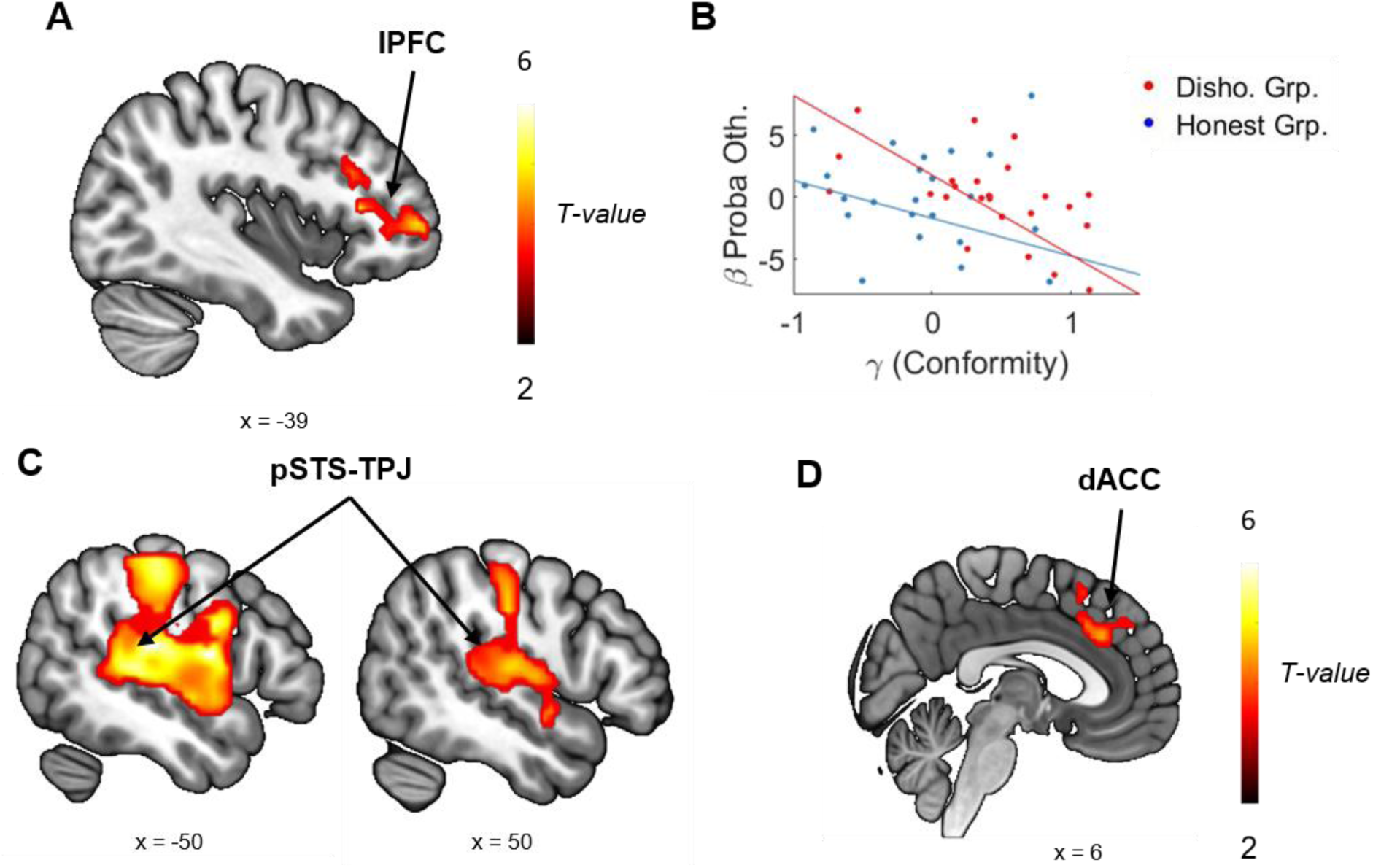
Neuroimaging results. **A**. Negative relationship between the BOLD signal in the left lateral prefrontal cortex (lPFC) and the dynamic valuation bias signal (i.e., the learned probability that the other cheats) with participants’ conformity parameter γ at the decision stage of the Solo trials of the Dishonest Group condition (p < 0.001 uncorrected and p < 0.05 whole-brain cluster-corrected family-wise error (FWE)). **B**. Individuals betas extracted from 6mm sphere centered on peak activity in the lPFC are negatively correlated with the participants’ conformity parameter γ in the Dishonest Group condition. For negative value of γ, participants show anti-conformism and the correlation between the Betas and the probability of cheating is positive. For positive γ value, that express conformity, the correlation is negative or null. In the Honest Group condition, this relationship is not significant. **C**. BOLD signal in the bilateral posterior superior temporal sulcus (pSTS) and the temporo-parietal junction (TPJ) correlates with the participants’ inference about others’ cheating probability for both group conditions at the prediction stage of the Predict trials (*p <* 0.001 uncorrected and *p <* 0.05 whole-brain cluster-corrected family-wise error (FWE)). **D**. BOLD signal in the dorsal anterior cingulate cortex (dACC) correlates negatively with the relative value of the participants’ (DV(Chosen) – DV(Unchosen)) in the Solo trials in the whole experiment (*p <* 0.001 uncorrected and *p <* 0.05 whole-brain cluster-corrected family-wise error (FWE)).

In the second GLM (GLM 2), we added the remaining three computational signals: the participants’ prediction that the other cheated at the time of the prediction in the Predict trials, their prediction error at the time of the feedback in the Predict trials and the relative value of the participants’ choice at the choice stage of the Solo trials in every condition (Baseline and Groups; see Methods SI for details). Using this GLM, we found that the prediction that the others cheat is encoded in the bilateral posterior Superior Temporal Sulcus and Temporo-Parietal Junction (pSTS-TPJ, x,y,z = 54,0,-9, −36, −34, 48; see Fig. 4.C and Table S4). Additionally, the relative value of the chosen option (𝐷𝑉_𝐶ℎ𝑜𝑜𝑠𝑒𝑛_ − 𝐷𝑉_𝑈𝑛𝑐ℎ𝑜𝑜𝑠𝑒𝑛_) in the Solo trials was negatively correlated with the BOLD signal in the dorsal Anterior Cingulate Cortex (dACC, x,y,z = 18, 27 48) regardless of blocks (i.e. blocks 1-3 averaged together) (Fig. 4.D and Table S5). Finally, the prediction error at the time of the feedback in the Predict trials, correlated with the bilateral ventral striatum (x,y,z = −12,3,-12; Fig. S3 and Table S6).

## Discussion

Individuals possess only a limited understanding of others’ opinions, intentions and preferences. Yet, social influence hinges on the ability to infer others’ likely actions. Our findings indicate that when we observe others’ cheating behavior, our brain monitors their inclinations toward dishonesty, not by shaping our own moral preferences, but by biasing our assessment of the value of cheating (valuation bias). Moreover, this valuation bias is dynamic. It reflects progressive learning about the moral inclinations of others. At the brain system level, participants who are anti-conformists (conformists) tend to show higher (lower) left lPFC activity with this valuation bias when observing dishonest behavior. Moreover, others’ dishonesty level is encoded in the TPJ-pSTS when participants predict their behavior. These results provide a dynamic and mechanistic understanding of social influence in cheating, and demonstrate the interplay between social learning and social influence processes.

At the behavioral level, participants changed their cheating behavior when they were exposed to the Dishonest Group, but not the Honest Group. This asymmetry in social influence could be driven by the difference in learning accuracy between the Dishonest and Honest groups. However, this interpretation is ruled out by the fact that participants discriminated the levels of dishonesty between the two groups. Moreover, the conformity parameter of the Dynamic Bias computational model was significantly higher when participants observed the behavior of the Dishonest Group than the Honest Group. This result echoes with previous research showing that anti-social behaviors such as cheating are more contagious than prosocial behavior or honesty (11, 30, 31). In addition, social proximity exacerbates this asymmetry (11) and can improve social learning (36). In our experiment, group members were anonymous. Thus, the asymmetry of influence cannot be explained by different levels of social proximity between the participants and others.

Two main mechanisms of social influence have been proposed: an adjustment in individuals’ preferences (8, 12, 37) or a valuation bias in individuals’ valuation process (3, 4). Previous work proposed that these two mechanisms are dependent on the accessibility of social information. Precisely, when individuals have a full access to others’ behavior or beliefs, the preference shifting mechanism would be more prevalent (38). Here, we show that a dynamic valuation bias better describes the mechanism of social influence when others’ behavior is fully accessible and accurately learned. Our model assumes that participants infer the preferences of others over time (social learning component). This internal representation then biases the valuation process of participants when they face the opportunity to cheat (social influence component). Such a process is in line with the proposal that social influence does not consist of blindly following others, nor of changing one’s preferences or the valuation process, but is rather the building of a causal understanding of others’ behavior, which then influences the decision process (28, 39, 40).

One original aspect of our experimental design is the way participants were gradually exposed to others’ behavior. Past experiments either presented others’ choices in a block of trials or did not consider the learning dynamics (4, 8). It should be noted that participants learned quickly, as they were able to accurately predict others’ behavior after approximately 10 trials. Despite this limitation, our dynamic model of social influence was able to explain participants behavior better, demonstrating that ‘fixed’ models are not the most representative of the process of social influence. One natural extension of our work would be to test a more volatile or uncertain social environment as this would induce a slower learning pace (41).

Our results elucidate the neurocomputational mechanisms of social influence. When exposed to a dishonest group, we observed differences in the dynamic valuation bias in the left lPFC, depending upon individual levels of conformity. That is, higher lPFC valuation bias activity was observed in participants who were more anti-conformist (i.e., resistant to the observed norm of cheating). This effect at the brain system level parallels the fact that the larger the valuation bias, the more likely participants were to cheat. Our observation of the dynamic tracking of the valuation bias by the dlPFC extends previous reports that the lPFC encodes the view of others about a choice, moral or not (8, 16). Furthermore, lPFC is known to be involved in norm compliance (21, 42). These previous results were obtained when the social norm was known from the beginning (i.e., not learned), unlike in our experiment. Here, we additionally show that the lPFC tracks the evolution of the social norm (i.e., the behavior of the others) depending on the participants’ degree of compliance. This shows that the lPFC tracks the social environment in a dynamic fashion and is responsible of the integration of social information into the decision valuation process.

In our Bayesian preferences learning model, based on the model developed by (35), individuals start with priors about the preferences of the others, which are then used to predict their likelihood to cheat. A prediction error updates those priors and corresponds to the difference between the predicated likelihood to cheat and the other’s actual behavior. During the feedback phase, this prediction error was encoded in the bilateral ventral striatum. This region has been demonstrated to encode prediction errors related to social norms, a process required for learning about the behavior of other individuals (43, 44). When investigating brain areas encoding the belief about the level of dishonesty of the group members, the bilateral pSTS and TPJ tracks one’s belief about the probability that a group member will cheat, when the participant is predicting her choice (both in the Dishonest and Honest Group conditions). Thus, the pSTS-TPJ computes a signal required for learning and predicting others’ behavioral tendencies. This is in line with previous work showing involvement of this brain region in simulating others’ learning or supporting the representation of others’ preferences (25, 45). Furthermore, we observed no implication of the pSTS-TPJ in the social influence process itself, as this was carried by the lPFC (i.e., which encoded the dynamic valuation bias). In contrast, most previous reports that the pSTS-TPJ is involved in the decision process, are based on experiments in which social information is directly useful for the participants to make a choice. This has been shown when choosing to cooperate (46), when achieving a consensus (25) or when revising one’s choice (47). In the current study, participants did not directly need social information to decide whether to cheat or not but could be influenced by what they learned about others’ behavior. Consistent with this, previous work on social influence did not observe engagement of the pSTS-TPJ when social learning is not involved (4, 8).

Overall, our neuroimaging results demonstrate that the pSTS-TPJ and dlPFC compute two separate signals: a social learning signal in the pSTS-TPJ and a social influence signal in the lPFC. Our study elucidates the dynamic link between social learning and social influence in the context of cheating behavior. Our computational model explains how individuals generate a formal representation of others’ preferences which then biases their valuation process. The representation of others’ behavioral tendencies was encoded in the rTPJ-pSTS region during learning while the dynamic bias engages the left lPFC. Our findings have implications for developing new computational theories of social influence that could be useful in many different contexts. For example, a recent fMRI study reported that cognitive processes underlying social understanding were more aligned after natural consensus-building conversation (48). Much research on social influence has focused on public compliance, setting aside the long-lasting effects of social interaction on private cognition (49, 50). In fact, consensus-building conversation not only align neural responses within groups (reflecting social compliance), but this alignment can generalize to novel stimuli that were not discussed (48). This emphasizes an important role of private acceptance (an effect explained here by the valuation bias) in understanding social influence.

## Methods

This study was approved by the local ethics committee (CPP Sud-Est 2, 2018-36). We provide a detailed description of the experimental procedures in SI Methods. In Solo trials, participants observed the outcome of die throw and had to report the result (Fig 2.A). They could either report the correct outcome or could cheat by reporting an incorrect outcome. Each option was associated with a payoff and cheating was always more rewarded than reporting honestly. In Predict trials, participants also observed the outcome of a die roll and were presented with two options, a, accurate and an inaccurate one (Fig. 2.A). However, they had to predict the option reported by another individual randomly selected from a group of 10 who performed the Solo trials in a previous experimental session. Unbeknownst to the participant, the group members were simulated. One group was dishonest and another was honest. The experiment was organized in three blocks (Fig. 2.B). A first block, the Baseline, was composed by 50 Solo trials. The purpose was to assess the initial preferences of the participants. The second and third blocks consisted of 50 Solo trials and 50 Predict trials that were interleaved starting with a Predict trial. Thus, participants were gradually exposed to others’ cheating behavior. The difference between these two blocks was the cheating behavior of the presented group (either dishonest or honest). The order of presentation of the dishonest and honest group was randomized between participants.

## Acknowledgements

This research has benefited from the financial support of IDEXLYON from Université de Lyon (project INDEPTH) within the Programme Investissements d’Avenir (ANR-16-IDEX-0005) and of the LABEX CORTEX (ANR-11-LABX-0042) of Université de Lyon, within the program Investissements d’Avenir (ANR-11-IDEX-007) operated by the French National Research Agency. This work was also supported by grants from the Agence Nationale pour la Recherche to JCD (ANR-21-CE37-0032), and by MITI 2020 CNRS to JCD. We would like to thanks Rémi Phillipe, Marie Devaine, Jean Daunizeau and Mateus Joffily for fruitful discussions to build our computational model, Franck Lamberton and Danielle Ibarrola from the Neuroimaging center CERMEP for the assistance in running the fMRI sessions.

## Supplementary Information

### Methods SI

#### Participants

32 healthy participants were recruited *via* the recruitment Facebook page of the Institute of Cognitive Sciences. Exclusion criteria included a history of systemic or neurological disorders, psychiatric disorders, psychoactive medication or drug use, pregnancy, involvement in psychology classes, and previous participation in studies involving decision-making in general. We recruited only right-handed participants. One participant was excluded from any analysis because they cheated in all the trials and reported that they did not believe the scenario, leaving 31 participants (14 males, mean age: 22.87 years, s.d. 0.58). For the model estimation and the fMRI results three additional participants were excluded because they never cheated in the Baseline condition. Thus, these analyses are made on 28 participants.

#### Experimental design

The task was based on a cheating game in which participants observed the outcome of a 6-sided die throw and had to report the result on a subsequent screen. Participants could either report the correct outcome (be honest) or could cheat by reporting a dishonest option. Each choice was associated with a payoff. To create a conflict between honesty and maximizing earnings, cheating was always the more profitable choice. Participants in our task performed a variation of this cheating game, in which we introduced two types of trials, the Solo trials and the Predict trials (Fig. 2.A and B).

##### Solo trials

In these trials, participants played the cheating game described above. After a jittered inter-trial interval (ITI, 3-7 s), the outcome of the 6-sided dice roll was presented for 1 s, then participants had an unlimited time to choose which outcome to report, the left or the right one. The choice was made by pressing a button box with their right hand (either the right or the left button). The honest and dishonest choice sides were randomly determined for each trial. After the button press the screen froze for 0.5 s. The task involved no cue to confirm the decision, in order to maximize the participants’ perception of unaccountability (17).

##### Predict trials

Predict trials commenced in a similar fashion to Solo trials because participants observed the outcome of die roll and were presented with two options, an accurate and an inaccurate one. However, participants were told they had to predict the result another individual reported in a previous experimental session. This other individual was said to be randomly selected from a group of 10 participants who performed the Solo trials in a previous experimental session. After indicating their prediction (left or right button press, depending on the predicted outcome), the predicted outcome was framed in red for a randomized duration (2 - 5s). Then, participants received feedback during which the “real” outcome reported by the other was framed in green. If this frame was overlayed with the participants’ prediction, their prediction was correct, otherwise it was incorrect. A correct prediction was rewarded with €2 and €0 otherwise.

##### Groups simulations

Unbeknownst to the participant, the group of 10 individuals were simulated. Specifically, we simulated two groups composed either of honest or dishonest individuals. We chose to tell to participants that they had to predict the decisions made by a group of individuals who performed the task in a previous experimental session to increase the likelihood of conformism towards the others’ behavior (51). To maintain the behavior of the “group members” homogeneous within each group, we actually constructed a dataset based on the responses of two participants that performed the online pilot of the task. Specifically, we selected them based on the congruence of their behavior (no erratic behavior) and their cheating level. One cheated in 85% and the other in 20% of the Solo trials in the Baseline test. We then determined which utility functions best explained the pilot participants’ behavior, which was the variable cost of cheating utility function (see the section Utility functions for details). We then used the estimated parameters to simulate the cheating behavior of the group members in each Predict trial. One of the groups was based on the dishonest participant’s parameters and the other on the honest participant’s parameters. This procedure ensured a high level of credibility as the decisions were simulated from the true behavior of two participants. Furthermore, using a simulation allows us to add noise to the data to decrease the pace of learning by our participants. This gave us two sets of Predict trials, one in which “others” were cheating in 22% of the trials (Honest group), the other in which the “others” cheated in 88% of the trials (Dishonest group, see Table S.1).

##### Task sequence

(Fig. 2.B). The experiment was divided into three blocks and was inspired by a previous experiment on social influence affecting risky choices (8). The first block, the Baseline, was composed by 50 Solo trials. The purpose of this block was to assess the initial preferences of the participants. The second and third blocks consisted of 50 Solo trials and 50 Predict trials that were interleaved starting with a Predict trial. The fact that participants were gradually exposed to the others’ (dis)honest behavior allowed us to test the potential link between learning about others’ behavior and the participants’ decisions to cheat. The difference between these two blocks was the cheating behavior of the group whose decisions the participants had to predict. One group was considered as dishonest (Dishonest Group condition, 88% cheating frequency) while the other was considered as honest (Honest Group condition, 22% cheating frequency). The order of presentation of the second and third blocks was randomized between participants. To prevent participants from simple imitative behavior, as they could simply repeat the behavior from the previous Predict trials, the parameters (payoffs and dice scores) of any Predict trial were never the same as the next three Solo trials.

##### Parametrization

The die values ranged from 1 to 6 and the payoffs ranged from €2 to €10 with €2 increments. Both the dice values and the payoffs were randomly selected with two constraints. First, we aimed to control for any correlation between the dice values and the payoffs. Consequently, specific dice values were not associated with fixed payoffs. Second, we aimed to have the same number of observations per value of relative earnings (difference between the payoff of the dishonest choice and the honest one). Based on the possible payoffs, 5 levels of relative earnings were possible, €2, €4, €6, €8 or €10. We randomly selected 5 payoff combinations (payoffs for the honest and dishonest reports) for each level of relative earnings, leading to a set of 25 trials (each with 2 die values and 2 payoffs). We then repeated this set twice for each block of the experiment, leading to a total of 50 trials whose order of presentation was randomized between participants.

#### Procedure

The study took place at the Neuroimaging Center CERMEP (https://www.cermep.fr/cermep_en.php) and was approved by the local ethics committee (CPP Sud-Est 2, 2018-36). During the medical screening, participants provided informed consent. Before entering the fMRI scanner, participants were asked to read privately the instructions of the task. A questionnaire was added at the end of the instructions to ensure they understood the task fully, and especially the fact that their decisions directly affected their final payment. We also made sure that participants were aware of the true anonymity of their decisions during the task. After the completion of the task, participants completed a debriefing questionnaire. They had to state on a 5 -point Likert scale (5 =” I completely believe it”, 1 =” I completely disbelieve it”) whether they believed that their decisions were kept confidential (mean = 4.42 ± 0.99), whether they believed their decision were scrutinized (mean = 3.39 ± 1.41) and whether they believed the others individuals were real (mean = 4.03 ± 1.02). These results confirmed that participants trusted that their decisions were confidential and that they believed that the group of others existed. To avoid a spillover effect on the cheating behavior in the task we also asked participants, at the end of the task, whether they thought it was immoral to cheat at the beginning of the task and at the end. The goal was to assess if the exposure to others’ dishonest behavior led to a change in their moral perception of cheating in our task. Ten participants reported, both for the beginning and end of the task, that it was never immoral to cheat, ten reported that it was immoral at the beginning of the task but not immoral at the end, seven reported that it was immoral, both for the beginning and end of the task, three reported that they had no opinion, at the beginning of task, but thought that it was not immoral at the end of the task. Finally, one participant had no opinion both at the beginning and the end of the task. Finally, we asked them if they had any comments or remarks about the task.

#### Statistical analysis

##### Behavioral analysis

Our behavioral statistical results are derived from mixed-effect logistic regressions. We always report the marginal effect rather than the odd ratios as it is easier to read and understand. The marginal effect can be interpreted as the mean discreet change of the dependent variable given a unitary change of an independent variable. In all regressions we considered standard errors clustered at the participants’ level as well as a random-effect at the participant level. When reporting pairwise comparisons of marginal effects we always controlled for multiple comparisons using a Bonferroni correction. Other statistical tests are always two-sided nonparametric tests unless specifically indicated otherwise.

##### Utility functions

Our different models are based on a core utility function derived from previous work in behavioral economics (33, 34). These models assume that agents compute a relative decision value of cheating by solving a cost-benefit arbitration between the relative gains of cheating and its moral cost (Δ𝐷𝑉 (𝑐ℎ𝑒𝑎𝑡)). Formally, the relative gains of cheating are weighted by a free parameter *α* representing the agent’s preference for money. The moral cost of cheating is a free parameter *δ* which is fixed (Fixed cost utility, see Eq. 1). We also considered an alternative utility function in which the cost of cheating weights the payoff of cheating (24)) (Variable cost utility, see Eq. 2).

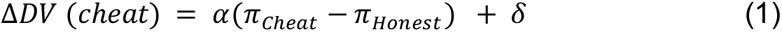

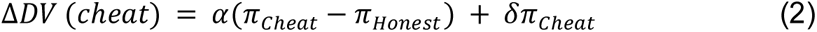

##### Learning models

To capture the computational processes underlying learning about others’ cheating behavior, we considered a family of 6 Bayesian learning models. All of them are based on the Bayesian Preferences Learning model (BPL) (35). In these models, participants infer others’ preferences from others’ observed choices. Formally, others’ preferences correspond to their own parameters *α* and *δ* for their utility function with this set defined as 𝛩(𝑜). This allows us to compute a relative decision value based on Eq. 1 or Eq. 2 which is then transformed to a probability using a softmax function with a temperature parameter *β* that is treated as a free parameter. Participants are assumed to start with prior beliefs about the others’ parameters 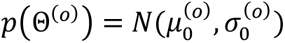 which are Gaussian with mean 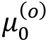 and variance 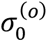. Given the others’ choices, the participants estimate and update the set of parameters 𝛩(𝑜) using the following Bayes-optimal probabilistic scheme:

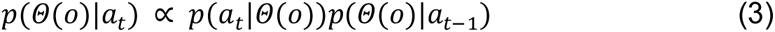

Where 𝑝(𝛩(𝑜)|𝑎_𝑡_) is the participant’s posterior belief about the other’s preference at the end of trial *t* and the right part of the equation represents the Bayesian belief update rule. Following (35), we used a variational-Laplace scheme to implement this model which yields 𝑝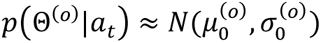. For more details about the formal mathematical description, we refer the interested reader to the original paper (35). As the BPL model assumes that participants start with priors about the others’ preferences, we considered 3 types of priors. In a first model, µ_𝛼_ and µ_𝛿_ were equal to the value of *α* and *δ* of the participant’s own utility function, estimated in the Baseline (BPL-Self). According to this model, participants are assumed to consider the group members as individuals with preferences similar to their own (self-projection). In a second model, we set the priors as in the previous case, but only for the first group they faced (Dishonest or Honest group conditions). For the second group, µ_𝛼_ and µ_𝛿_ were equal to µ_𝛼_ and µ_𝛿_ learned at the last Predict trial of the first group they faced (BPL-Others). Here, participants are supposedly projecting the learned preferences of the first group onto the second one. In a third and last model, the setting was not changed for the first group, but only for the second, µ_𝛼_ and µ_𝛿_ were the weighted means of the participant’s parameter values (as in the BPL-Self) and of the value learned for the previous group (BPL-SelfOthers). This weight is a free parameter 𝜔.

Finally, we considered either one of the two utility functions to be the one used by our participants to infer the others’ preferences. In total, we tested the 3 different types of prior for each utility function (see Eq. 1 and Eq. 2) leaving us with 6 candidate models.

##### Social influence models

Our computational models to account for social influence are built on two potential mechanisms. The first one, the Preferences Shift (PS), assumes that participants’ preferences change when they are exposed to others’ cheating behavior (8, 12). The second one, the Valuation Bias (VB), assumes that participants’ decision values change, while their preferences remain constant, when they are exposed to others’ cheating behavior (3, 4, 18). We derived formal computational models from these two mechanisms. For the Fixed Preferences Shift (PS Fixed) model this is formally defined as follows:

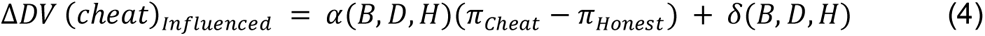

In this equation (Eq. 4), the participants’ preferences *α* and *δ* take different values for each condition (Baseline, Dishonest and Honest Groups: *B,D,H*). For the Fixed Valuation Bias (VB Fixed) mechanism this is formally defined as follows,

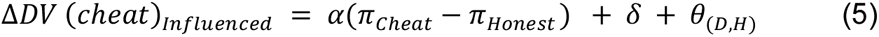

In this equation (Eq. 5), the participants’ preferences are held fixed throughout the task. A parameter θ represents the bias of the participants’ decision values due to the exposition to others’ cheating behavior. This parameter is different for the Dishonest Group (θ_𝐷_) and for the Honest Group (θ_𝐻_).

In addition to these two models, we also considered dynamic variants that imply that the preferences change or the value biases are affected by the participants’ knowledge of the others’ cheating behavior. The main idea is that, over observations of others’ behavior, our participants learn about the preferences of others’. Over the course of this learning, either the changes in our participants’ preferences or the valuation bias evolved as a result of their observations. Formally, we tested 2 different models that link the outcome of the BPL model with either the PS (Eq. 4) or the VB model (Eq. 5).

In the first model, we considered that the changes of preferences are the results of a weighted average of the participants’ own preferences and what they know at a given point in time about the others’ preferences. Formally, the Dynamic Preference Shift (PS Dynamic) model is defined as follows,

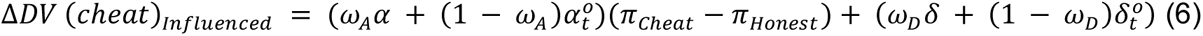

where 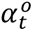 and 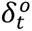 are the others’ preferences derived from the BPL model at time *t*, 𝜔*_A_* ( 𝜔*_D_*) being free parameters representing the magnitude of social influence. The closer from 0 these parameters are, the more participants will be influenced by their perception of the others’ preferences, i.e., the more they will be susceptible to social influence (with 0 ≤ 𝜔_𝐴_(𝜔_𝐷_) ≤ 1). Formally, they control the weight that participants put on their own preference parameter *α* (*δ*) and 1 − 𝜔_𝐴_ (1 − 𝜔_𝐷_) is the weight that participants put on others’ learned preferences *α_t_^O^* (*δ_t_^O^*). Both 𝜔_𝐴_ and 𝜔_𝐷_ were estimated separately for the Dishonest and Honest Group conditions.

In the second model, we considered that the value bias is derived from what participants have learned about others’ preferences at a given point in time Formally, the Dynamic Valuation Bias (VB Dynamic) model is defined as follows,

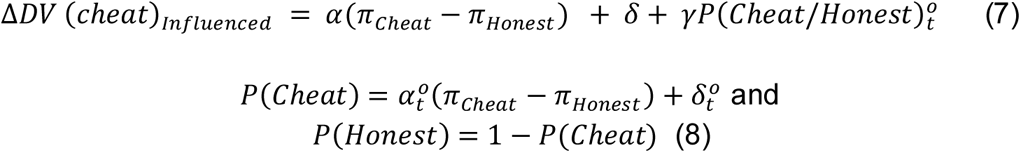

where 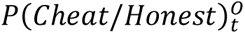 being the dynamic valuation bias. It corresponds to the probability that others’ cheated (were honest) in the participant’s position. It is based on what participants learned about others’ preferences (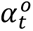 𝑎𝑛𝑑 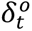) at time *t* in each of the two group conditions and on the relative payoff of the given Solo trials. *γ* is a free parameter representing the extent of the participants’ conformity. This last parameter is estimated separately for the Dishonest Group and the Honest Group conditions.

Overall, we tested 4 model candidates to account for social influence in our experiment. Participants were assumed to use the Fixed Moral cost utility function to compute their decision value as it is the utility function that best explained our participants’ behavior throughout the experiment (see Fig. S1.A and the Results section for details).

##### Model selection

The Bayesian Model Selection (BMS) was performed using the VBA toolbox (Variational Bayesian Analysis) in a random effect analysis relying on the free energy as the lower bound of model evidence. We use protected Exceedance Probability measurements (pEP) to select the model used most frequently in our population of participants (32).

#### fMRI acquisition and preprocessing

MRI acquisitions were performed on a 3 Tesla scanner using EPI BOLD sequences and T1 sequences at high resolution. Scans were performed in a Siemens Magnetom Prisma scanner HealthCare at CERMEP Bron (single-shot EPI, TR / TE = 1600/30, flip angle 75, multiband acquisition (accelerator factor of 2), in an ascending interleaved manner with slices interlaced 2.40 mm thickness, FOV = 210 mm. We also used the iPAT mode with an accelerator factor of 2 and the GRAPPA method reconstruction. The number of volumes acquired varied given the time the participant took to make their decisions. The first acquisition was made after stabilization of the signal (3 TR). Whole-brain high-resolution T1-weighted structural scans (0.8 x 0.8 x 0.8 mm) were acquired for each subject, co-registered with their mean EPI images and averaged across subjects to permit anatomical localization of functional activations at the group level. Field map scans were acquired to obtain magnetization values that were used to correct for field inhomogeneity.

#### fMRI data analysis

Image analysis was performed using SPM12 (Wellcome Department of Imaging Neuroscience, Institute of Neurology, London, UK, fil.ion.ucl.ac.uk/spm/software/spm12/). Time-series images were registered in a 3D space to minimize any effect that could result from participant head-motion. Once DICOMs were imported, functional scans were realigned to the first volume, corrected for slice timing and unwarped to correct for geometric distortions. Inhomogeneous distortions-related correction maps were created using the phase of non-EPI gradient echo images measured at two echo times (5.20 ms for the first echo and 7.66 ms for the second). Finally, in order to perform group and individual comparisons, they were co-registered with structural maps and spatially normalized into the standard Montreal Neurological Institute (MNI) atlas space using the DARTEL method.

We ran general linear models (GLMs) analysis to identify which brain regions encoded: (1) the dynamic valuation bias in both the Dishonest and Honest Group conditions (choice stage in the Solo trials); (2) the learned probability that the others cheated, in the Predict trials in both the Dishonest and Honest group conditions (prediction stage in the Predict trials); (3) the relative value of participants’ chosen options in Solo trials, regardless of conditions. We also included a fourth signal, the prediction error at the feedback stage of the Predict trials. In every GLM an event was defined as a boxcar function whose duration was equal to the participant’s reaction time or to the time of display with the exception of the button press which was defined as a stick function. Events such as the stimulus stage, the choice and prediction stages for both Solo and Predict trials as well as the feedback stage in the Predict trials were always included in all of our GLMs. Head movement parameters were added as parametric regressors of no interest to account for motion related noise. Finally, to control for diverse task parameters, we added as parametric regressors in every GLMs for the choice and prediction onsets: the choice or prediction of the participant (0: No cheat, 1: cheat), the participants’ reaction time, and which side was pressed in the choice or prediction stage of the Solo and Predict trials, respectively. Based on these common features we defined 2 GLMs.

In GLM1, we added the following parametric regressors: the inferred others’ probability to cheat (be honest) in the choice stage of the Solo trials of the Dishonest (Honest) group condition (dynamic valuation bias, signal 1), as well as the relative value of cheating derived only from participants’ preferences also at the choice stage of the Solo trials but in every condition. This last parametric regressor represents the unbiased participants’ relative valuation of cheating. In GLM2, we added the following parametric regressors: the learned probability that the others’ cheated at the prediction stage of Predict trials in both the Dishonest and Honest Group conditions (signal 2). In the choice stage of the Solo trials, we added the relative value of the participants’ choice in every condition (signal 3). Then, in the feedback stage of the Predict trials we added the prediction error from the BPL-Self model in both the Dishonest and Honest Group conditions. Finally, in both GLMs, we added the participants’ conformity parameters *γ* as a second-level covariate, for the Dishonest and Honest Group conditions. We computed one sample t-tests with contrasts for main effect for each parametric regressor as well as the main effect per block (Baseline, Dishonest and Honest groups) when possible. Reported brain areas show a significant activity at the threshold of *p <* 0.05, whole brain family-wise error (FWE), corrected for multiple comparisons at the cluster level (threshold at *p <* 0.001 uncorrected).

## Figures

**Figure S1:**
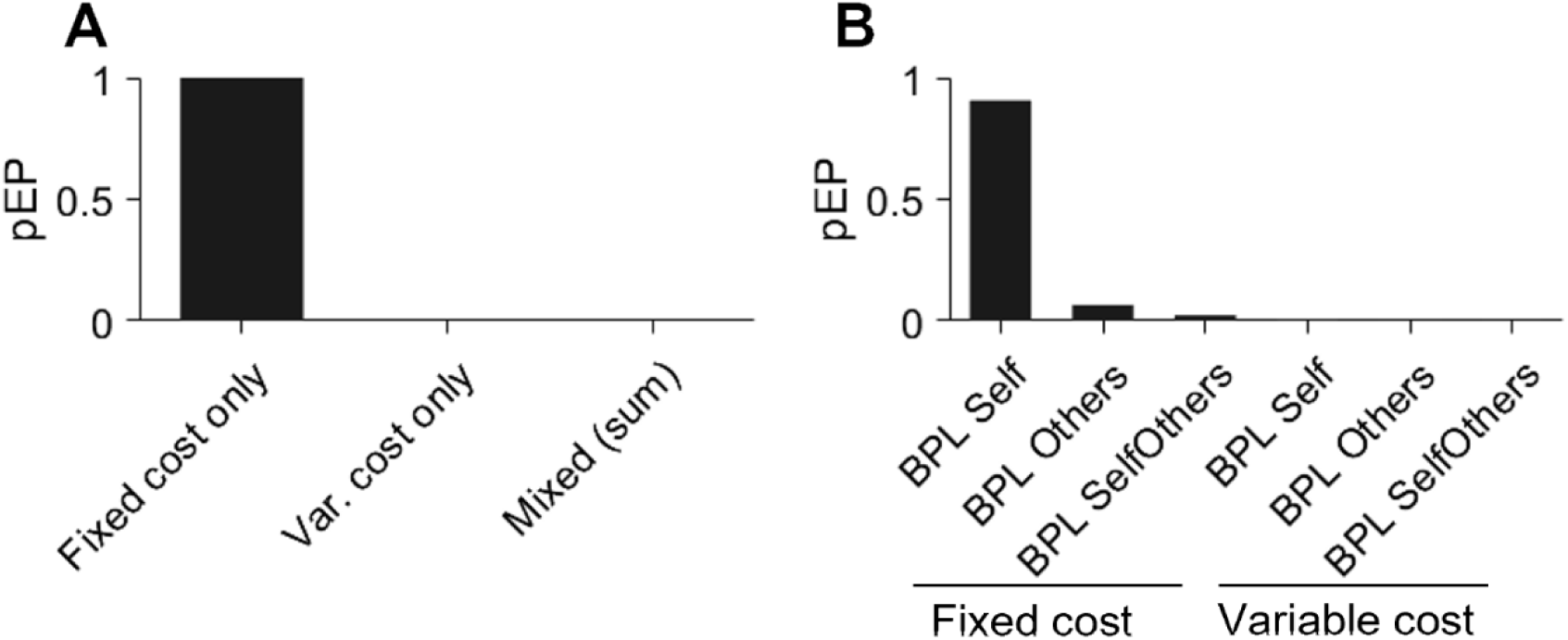
Utility function and social learning model selection. **A.** Protected Exceedance Probability (pEP) of the different utility functions associated with cheating, estimated over the three blocks of the experiment. The higher the pEP the more likely a given model explains the group’s behavior more than the others. The fixed cost only utility function assumes that participants are bearing a fixed moral cost when choosing to cheat while the variable cost variant assumes that this moral cost depends on the cheating payoff. The mixed variants are all the possible combinations of these two utility functions across the three blocks of the experiment (baseline and groups conditions). **B.** Protected Exceedance Probability (pEP) of the different social learning models. The higher the pEP the more likely a given model explains the group’s behavior more than the others. We tested a total of 8 models all based on the Bayesian Preference Learning model (BPL) for which we varied the priors concerning the others’ parameters (Self, Others or SelfOthers) and whether the utility function the participants used to learn about the others’ cheating behavior was the fixed cost or the variable cost utility function.

**Figure S2:**
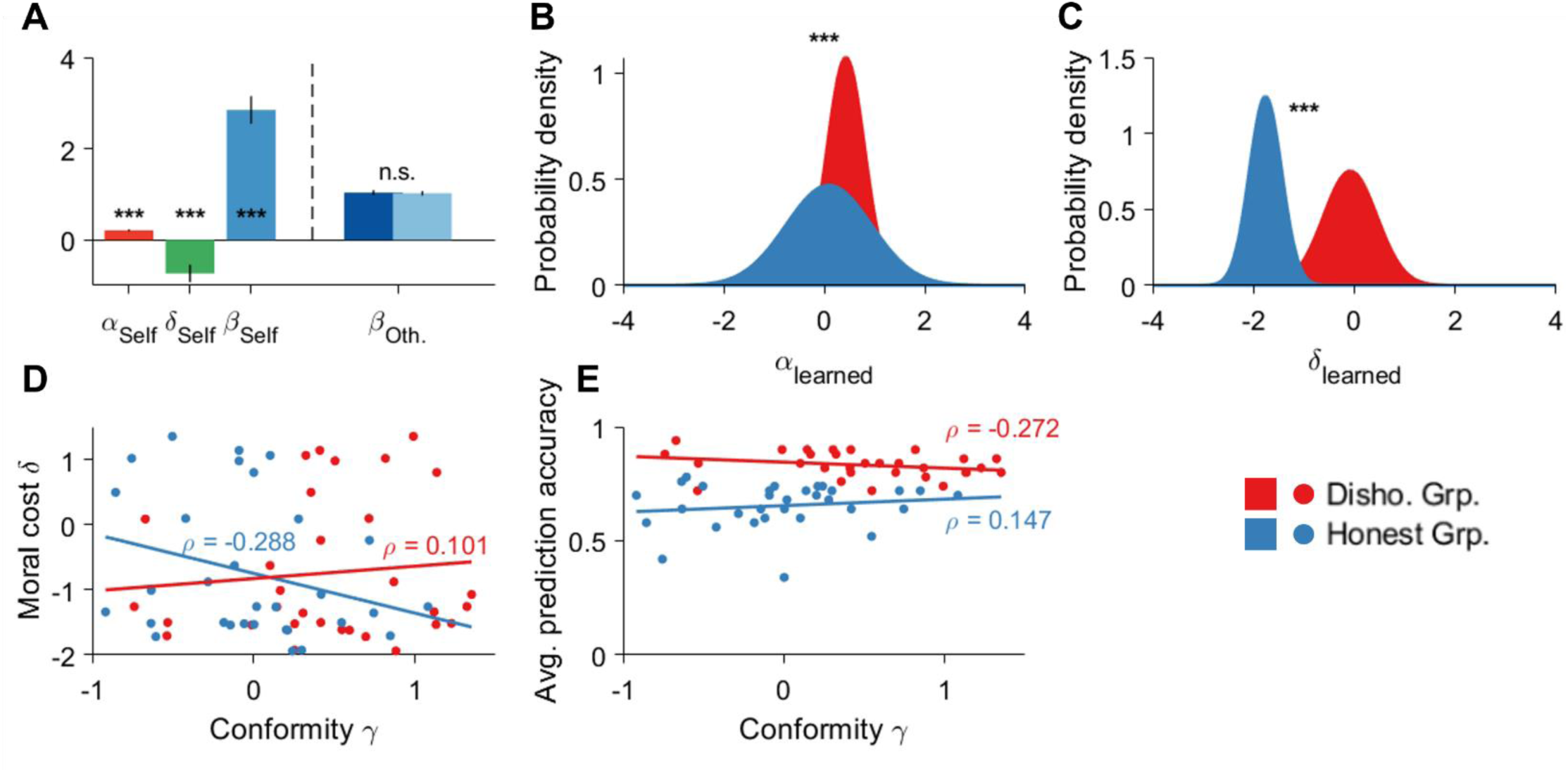
Model selection and estimation. **A.** Mean values of the participants’ social influence model parameters (left side) and the learned temperature for the Dishonest Group and Honest Group condition (right side). Statistical tests for the influence model parameters correspond to one-sided signed rank tests against 0 and for the learned temperature it corresponds to two-sided rank-sum tests comparing the values between the Dishonest and Honest group blocks (in dark and light blue, respectively). Bars correspond to standard errors. **B-C.** Probability density function of the learned parameters *α* (**B**) and *δ* (**C**) for both the Dishonest and Honest Group conditions. Stars correspond to the p-values of a two-sided rank-sum test comparing the mode of distribution between the two conditions. **D**. The participants’ moral cost parameter 𝛿 is not correlated with the participants’ conformity parameter 𝛾 in either the Dishonest or Honest Group conditions. The values 𝜌 are the correlation parameters obtained from a Pearson correlation (p=0.588 and p=0.116 for the Dishonest Group and Honest Group conditions, respectively). **E.** The participants’ average prediction accuracy is not correlated with the participants’ conformity parameter γ in either the Dishonest or Honest Group conditions. The values ρ are the correlation parameters obtained from a Pearson correlation (p=0.139 and p=0.431 for the Dishonest and Honest Group conditions, respectively). *** *p <* 0.001.

**Figure S3:**
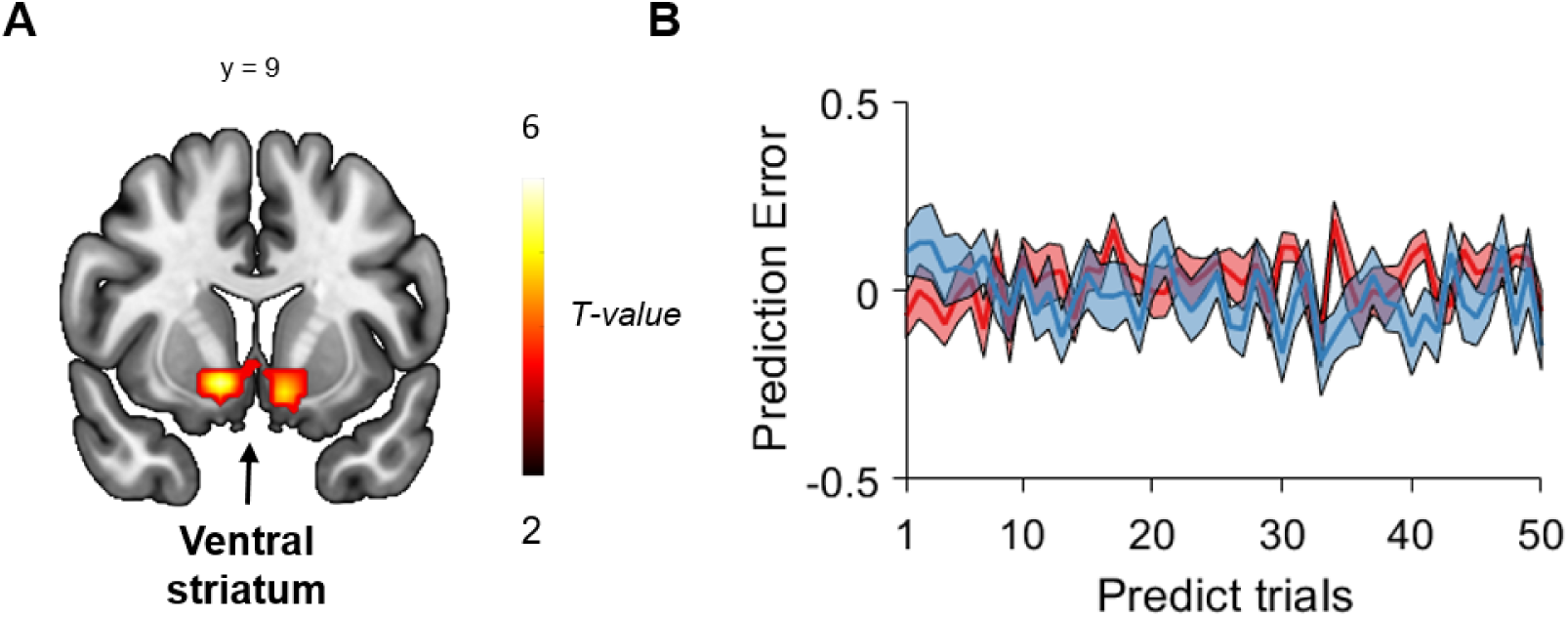
**A.** Regions encoding the prediction error from the social learning model at the time of the feedback in the Predict trials. FWE clustered-corrected with *p <* 0.05. **B.** Mean prediction error over predict trials for the Dishonest Group condition (red), and Honest Group condition (blue). Shaded areas are standard errors.

**Table S1:**
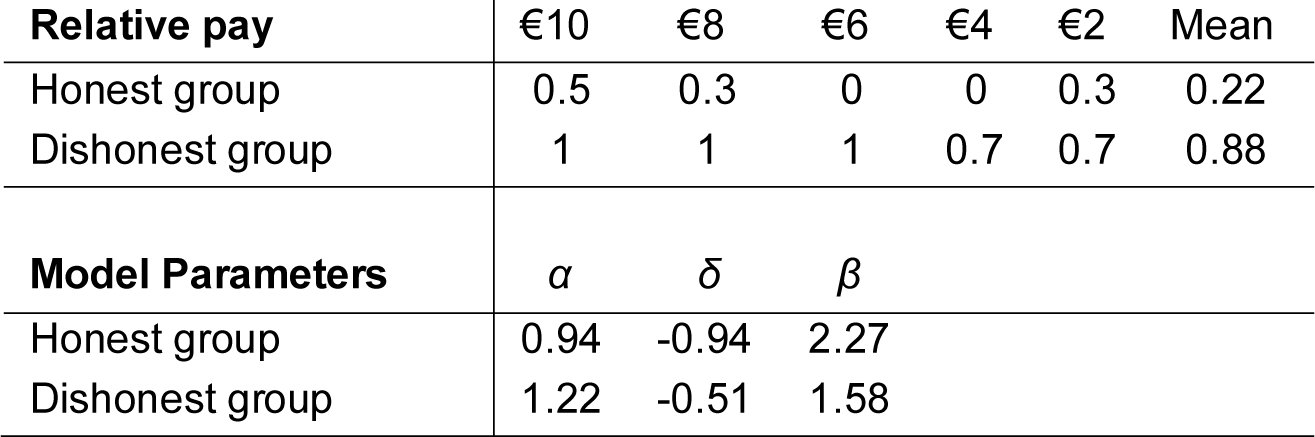
Mean cheating frequency per level of relative pay for cheating for the two groups and model parameters used to simulate the groups’ behavior (Variable Moral cost utility function).

**Table S2:** Logistic random-effect regressions.

|  | (1) | (2) | (3) |
| --- | --- | --- | --- |
|  | Lying<br>(1: Cheat, 0: No cheat) | Prediction accuracy<br>(1: Correct pred., 0: Incorrect pred.) | Prediction lying<br>(1: Lie pred, 0: No lie pred) |
| Disho. Grp. vs Baseline | 0.148 ***<br>(0.037) | -<br>- | -<br>- |
| Honest Grp. vs Baseline | 0.047<br>(0.025) | -<br>- | -<br>- |
| Disho. Grp vs Hon. Grp. | 0.101 *<br>(0.027) | 0.179 ***<br>(0.022) | 0.540 ***<br>(0.033) |
| Honest Grp. first | -0.067<br>(0.100) | -0.045 **<br>(0.018) | 0.034<br>(0.035) |
| Trial number | < 0.001<br>(< 0.001) | 0.002 ***<br>(< 0.001) | (< 0.001)<br>(< 0.001) |
| Relative payoff cheating | 0.049 ***<br>(0.006) | 0.010 ***<br>(0.002) | 0.049 ***<br>(0.004) |
| Diff. dice value | -0.013 **<br>(0.004) | < -0.001<br>(0.004) | -0.004<br>(0.004) |
| Demographics | Yes | Yes | Yes |
| Number of observations | 4650 | 3100 | 3100 |
| Number of clusters | 31 | 31 | 31 |
| $P > \chi^2$ | < 0.001 | < 0.001 | < 0.001 |

Notes: Relative payoff for cheating: $\pi_{\text{Cheat}} - \pi_{\text{Honest}}$ , Diff. dice value: $\text{Dice value}_{\text{Cheat}} - \text{Dice value}_{\text{Honest}}$ . Standard errors clustered at the participant level are in parentheses. \*\*\* p<0.001, \*\* p<0.01, \* p<0.05.

**Table S3:**
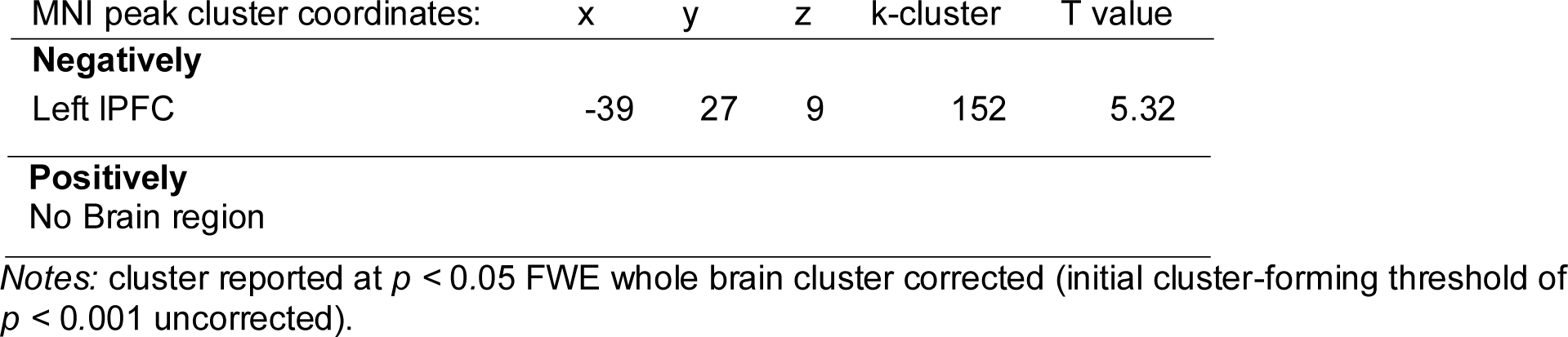
Brain regions encoding the Dynamic Valuation Bias in the Solo trials of the Dishonest Group condition modulated by the participants’ conformity parameter *γ*.

**Table S4:** Brain regions encoding the participants’ inferred probability that the other cheated at the time of the prediction in the Predict trials.

| MNI peak cluster coordinates: | x | y | z | k-cluster | T value |
| --- | --- | --- | --- | --- | --- |
| **Positively** |  |  |  |  |  |
| Right Superior Temporal sulcus | 54 | 0 | -9 | 1138 | 4.39 |
| Cerebellum | 30 | -45 | -33 | 197 | 4.76 |
|  | 15 | -66 | -57 | 509 | 4.54 |
| Left motor cortex | -36 | -24 | 48 | 3618 | 6.81 |
| Cuneus | 12 | -78 | 24 | 184 | 4.97 |
| **Negatively** |  |  |  |  |  |
| No Brain region |  |  |  |  |  |

Notes: cluster reported at $p < 0.05$ FWE whole brain cluster corrected (initial cluster-forming threshold of $p < 0.001$ uncorrected).

**Table S5:** Brain regions encoding the relative value of the participants’ choice in the Solo trials.

| MNI peak cluster coordinates: | x | y | z | k-cluster | T value |
| --- | --- | --- | --- | --- | --- |
| **Negatively** |  |  |  |  |  |
| dACC | 18 | 27 | 48 | 212 | 4.12 |
| **Positively** |  |  |  |  |  |
| No Brain region |  |  |  |  |  |

Notes: cluster reported at $p < 0.05$ FWE whole brain cluster corrected (initial cluster-forming threshold of $p < 0.001$ uncorrected).

**Table S6:** Brain regions encoding the prediction error at the time of the feedback in the Predict trials.

| MNI peak cluster coordinates: | x | y | z | k-cluster | T value |
| --- | --- | --- | --- | --- | --- |
| **Positively** |  |  |  |  |  |
| Ventral striatum | -12 | 3 | -12 | 177 | 5.93 |
| Left motor cortex | -30 | -18 | 51 | 207 | 3.68 |
| **Negatively** |  |  |  |  |  |
| dACC | 0 | 33 | 39 | 289 | 4.41 |
| Right angular | 30 | -54 | 42 | 210 | 4.37 |

Notes: cluster reported at $p < 0.05$ FWE whole brain cluster corrected (initial cluster-forming threshold of $p < 0.001$ uncorrected).

## References

1. F. Gino, S. Ayal, D. Ariely, Contagion and differentiation in unethical behavior: the effect of one bad apple on the barrel. Psychol Sci 20, 393–8 (2009).

2. J. Benistant, F. Galeotti, M. C. Villeval, Competition, information, and the erosion of morals. J Econ Behav Organ 204, 148– 163 (2022).

3. D. Chung, M. A. Orloff, N. Lauharatanahirun, P. H. Chiu, B. King-Casas, Valuation of peers’ safe choices is associated with substance-naïveté in adolescents. Proceedings of the National Academy of Sciences 117, 31729–31737 (2020).

4. D. Chung, G. I. Christopoulos, B. King-Casas, S. B. Ball, P. H. Chiu, Social signals of safety and risk confer utility and have asymmetric effects on observers’ choices. Nat Neurosci 18, 912–916 (2015).

5. M. A. J. Apps, N. Ramnani, Contributions of the medial prefrontal cortex to social influence in economic decision-making. Cerebral Cortex 27, 4635–4648 (2017).

6. M. Moutoussis, R. J. Dolan, P. Dayan, How People Use Social Information to Find out What to Want in the Paradigmatic Case of Inter-temporal Preferences. PLoS Comput Biol 12 (2016).

7. C. Calluso, A. Tosoni, G. Fortunato, G. Committeri, Can you change my preferences? Effect of social influence on intertemporal choice behavior. Behavioural Brain Research 330, 78–84 (2017).

8. S. Suzuki, E. L. S. Jensen, P. Bossaerts, J. P. O’Doherty, Behavioral contagion during learning about another agent’s risk-preferences acts on the neural representation of decision-risk. Proc Natl Acad Sci U S A 113, 3755–3760 (2016).

9. M. Fabbri, E. Carbonara, Social influence on third-party punishment: An experiment. J Econ Psychol 62, 204–230 (2017).

10. O. FeldmanHall, R. A. Otto, E. A. Phelps, Learning moral values: another’s desire to punish enhances one’s own punitive behavior. J Exp Psychol Gen 147, 1211–1224 (2018).

11. E. Dimant, Contagion of pro- and anti-social behavior among peers and the role of social proximity. J Econ Psychol 73, 66– 88 (2019).

12. H. Yu, J. Z. Siegel, J. A. Clithero, M. J. Crockett, How peer influence shapes value computation in moral decision-making. Cognition 211, 104641 (2021).

13. C. Qu, J. Bénistant, J. C. Dreher, Neurocomputational mechanisms engaged in moral choices and moral learning. Neurosci Biobehav Rev 132, 50–60 (2022).

14. G. Ugazio, et al., Neuro-Computational Foundations of Moral Preferences. Soc Cogn Affect Neurosci 17, 253–265 (2022).

15. M. A. Maréchal, A. Cohn, G. Ugazio, C. C. Ruff, Increasing honesty in humans with noninvasive brain stimulation. Proceedings of the National Academy of Sciences 114, 4360–4364 (2017).

16. M. J. Crockett, J. Z. Siegel, Z. Kurth-Nelson, P. Dayan, R. J. Dolan, Moral transgressions corrupt neural representations of value. Nat Neurosci 1–10 (2017). 10.1038/nn.4557.

17. C. Qu, E. Météreau, L. Butera, M. C. Villeval, J. C. Dreher, Neurocomputational mechanisms at play when weighing concerns for extrinsic rewards, moral values, and social image. PLoS Biol 17, e3000283 (2019).

18. G. D. A. Brown, S. Lewandowsky, Z. Huang, Social Sampling and Expressed Attitudes: Authenticity Preference and Social Extremeness Aversion Lead to Social Norm Effects and Polarization. Psychol Rev (2022). 10.1037/rev0000342.supp.

19. J. W. Buckholtz, Social norms, self-control, and the value of antisocial behavior. Curr Opin Behav Sci 3, 122–129 (2015).

20. R. W. Carlson, M. J. Crockett, The lateral prefrontal cortex and moral goal pursuit. Curr Opin Psychol 24, 77–82 (2018).

21. C. C. Ruff, G. Ugazio, E. Fehr, Changing social norm compliance with noninvasive brain stimulation. Science 342, 482–4 (2013).

22. A. Dogan, et al., Prefrontal connections express individual differences in intrinsic resistance to trading off honesty values against economic benefits. Sci Rep 6, 33263 (2016).

23. J. D. Greene, J. M. Paxton, Patterns of neural activity associated with honest and dishonest moral decisions. Proc Natl Acad Sci U S A 106, 12506–11 (2009).

24. L. Zhu, et al., Damage to dorsolateral prefrontal cortex affects tradeoffs between honesty and self-interest. Nat Neurosci 17, 1319–1321 (2014).

25. S. Suzuki, R. Adachi, S. Dunne, P. Bossaerts, J. P. O’Doherty, Neural mechanisms underlying human consensus decision-making. Neuron 86, 591–602 (2015).

26. T. E. J. Behrens, L. T. Hunt, M. W. Woolrich, M. F. S. Rushworth, Associative learning of social value. Nature 456, 245–249 (2008).

27. E. D. Boorman, J. P. O’Doherty, R. Adolphs, A. Rangel, The behavioral and neural mechanisms underlying the tracking of expertise. Neuron 80, 1558–1571 (2013).

28. C. J. Charpentier, K. Iigaya, J. P. O’Doherty, A Neuro-computational Account of Arbitration between Choice Imitation and Goal Emulation during Human Observational Learning. Neuron 106, 687–699.e7 (2020).

29. L. Zhang, J. Gläscher, A brain network supporting social influences in human decision-making. Sci Adv 6, 1–20 (2020).

30. P. Colzani, G. Michailidou, L. Santos-Pinto, Experimental evidence on the transmission of honesty and dishonesty: A stairway to heaven and a highway to hell. Econ Lett 231 (2023).

31. C. Bicchieri, E. Dimant, S. Gächter, D. Nosenzo, Social proximity and the erosion of norm compliance. Games Econ Behav 132, 59–72 (2022).

32. L. Rigoux, K. E. Stephan, K. J. Friston, J. Daunizeau, Bayesian model selection for group studies - revisited. Neuroimage 84, 971–985 (2014).

33. J. Abeler, D. Nosenzo, C. Raymond, Preferences for Truth-Telling. Econometrica 87, 1115–1153 (2019).

34. U. Gneezy, A. Kajackaite, J. Sobel, Lying Aversion and the Size of the Lie. American Economic Review 108, 419–153 (2018).

35. M. Devaine, J. Daunizeau, Learning about and from others’ prudence, impatience or laziness: The computational bases of attitude alignment. PLoS Comput Biol 13, 1–28 (2017).

36. W. Zou, X. Xu, Ingroup bias in a social learning experiment. Exp Econ 26, 27–54 (2023).

37. M. M. Garvert, M. Moutoussis, Z. Kurth-Nelson, T. E. J. Behrens, R. J. Dolan, Learning-Induced plasticity in medial prefrontal cortex predicts preference malleability. Neuron 85, 418–428 (2015).

38. H. Lee, D. Chung, Characterization of the Core Determinants of Social Influence From a Computational and Cognitive Perspective. Front Psychiatry [Preprint] (2022).

39. F. Cushman, Computational Social Psychology. Annual Review of Psychology 75, 625–652 (2024).

40. A. Najar, E. Bonnet, B. Bahrami, S. Palminteri, The actions of others act as a pseudo-reward to drive imitation in the context of social reinforcement learning. PLoS Biol 18 (2020).

41. T. E. J. Behrens, M. W. Woolrich, M. E. Walton, M. F. S. Rushworth, Learning the value of information in an uncertain world. Nat Neurosci 10, 1214–1221 (2007).

42. J. Gross, F. Emmerling, A. Vostroknutov, A. T. Sack, Manipulation of pro-sociality and rule-following with non-invasive brain stimulation. Sci Rep 8 (2018).

43. V. Klucharev, K. Hytönen, M. Rijpkema, A. Smidts, G. Fernández, Reinforcement Learning Signal Predicts Social Conformity. Neuron 61, 140–151 (2009).

44. T. Xiang, T. Lohrenz, P. R. Montague, Computational Substrates of Norms and Their Violations during Social Exchange. Journal of Neuroscience 33, 1099–1108 (2013).

45. S. Suzuki, et al., Learning to Simulate Others’ Decisions. Neuron 74, 1125–1137 (2012).

46. S. A. Park, M. Sestito, E. D. Boorman, J. C. Dreher, Neural computations underlying strategic social decision-making in groups. Nat Commun 10, 1–12 (2019).

47. A. Mahmoodi, H. Nili, D. Bang, C. Mehring, B. Bahrami, Distinct neurocomputational mechanisms support informational and socially normative conformity. PLoS Biol 20, 1–21 (2022).

48. B. Sievers, C. Welker, U. Hasson, A. M. Kleinbaum, T. Wheatley, Consensus-building conversation leads to neural alignment. Nat Commun 15, 3936 (2024).

49. A. G. Huth, W. A. De Heer, T. L. Griffiths, F. E. Theunissen, J. L. Gallant, Natural speech reveals the semantic maps that tile human cerebral cortex. Nature 532, 453–458 (2016).

50. U. Hasson, R. Malach, D. J. Heeger, Reliability of cortical activity during natural stimulation. Trends Cogn Sci [Preprint] (2010).

## Reference

51. Park, S. A., Goïame, S., O’Connor, D. A., & Dreher, J. C. (2017). Integration of individual and social information for decision-making in groups of different sizes. PLoS Biology, 15(6), 1–28.

